# Post-translational regulation of Platelet-derived growth factor receptor β is critical for fracture repair in aged mice

**DOI:** 10.1101/2023.11.04.565143

**Authors:** Jianguo Tao, Brea Lipe, Brendan F. Boyce, Lianping Xing, Hengwei Zhang

## Abstract

Increased protein ubiquitination was observed in fracture callus and particularly in aged mice. Treatment of proteasome inhibitor enhanced fracture repair in young and mice by increasing the number of mesenchymal progenitor cells (MPCs). However, the protein targets of proteasome inhibitor are still not known. Ub-proteomics identifies the top ub-proteins in MPCs and osteoblasts. Among them, PDGFRβ plays important roles both in osteogenesis and angiogenesis, which were reduced in callus of aged mice. We examine the dramatic decrease of PDGFRβ protein level, increased Ub-PDGFRβ, but mild decrease of mRNA in callus of aged mice, suggesting dys-regulated protein modification is the major cause of decreased PDGFRβ level. Decreased PDGFRβ results in the failure of PDGF-BB enhanced MPCs proliferation and fracture repair in aged mice. Co-treatment with proteasome inhibitor rescues the ability of PDGF-BB on MPC proliferation and fracture repair. Our findings not only discover the protein target of proteasome inhibitor in MPCs, but importantly connect the compromised effect of PDGF treatment on diseases with PDGFRβ proteasomal degradation. We open a new avenue for the treatment of fracture repair in elderly with the combination of PDGF-BB and proteasome inhibitor.

## Introduction

Ubiquitin-proteasome system (UPS) is an important cellular machine for protein post-translational modifications, which specifically adds to or removes ubiquitin from/to the substrate protein, resulting in degradation of ubiquitinated (Ub) substrate in proteasome or lysosomes. The substrate ubiquitination is carried out via sequential enzymatic reactions involving ubiquitin activating enzyme E1, ubiquitin-conjugating enzyme E2, and ubiquitin ligase E3 that confer substrate specificity by linking ubiquitin to target molecules^1^. A number of osteoblast (OB) positive regulators such as BMP^2^ and TGFβ^3^ signaling proteins, Runx2^4^(ref), and JunB^5^ are regulated via the UPS.

Bone fracture repair is composed of a series of events involving inflammation, angiogenesis, osteogenesis and remodeling. It was reported that aged mice have decreased mesenchymal progenitor cells (MPCs) and angiogenesis, which both are critical for fracture repair^6^. We confirmed their decrease in current study. We previously reported that CD45-Sca1+CD105+ MPCs^7^ from aged mice have increased UPS-mediated proteasome degradation of Runx2 and JunB proteins^5^.

Bortezomib (Btz) is FDA approved for treatment of multiple myeloma. It acts by inhibiting the proteasome via reversible occupation of the active proteolytic site of the 20S proteasome. It induces apoptosis in myeloma cells after the accumulation of excessive proteins^8^. In bone cells, Btz promotes OB differentiation by inhibiting Runx2 degradation^5^, and inhibits osteoclast (OC) formation by inhibiting the NF-kB pathway^9^. Btz and other proteasome inhibitors are very attractive candidates for the development of bone anabolic agents. Several studies performed in young or adult mice reported that Btz increased bone volume in normal and OVX mice^10^, and promoted fracture healing^11^. We previously reported that Btz enhanced fracture repair in adult young mice by markedly increasing Nestin+ MPC cells using Nestin-GFP mice^12^. Because aged mice have increased total ubiquitinated proteins in long bone and callus than young mice, we further studied the effect of Btz on aged mice and found Btz enhanced fracture repair in aged mice by increasing MPC number and angiogenesis^13^. Our findings suggested that proteasome inhibition may be a useful strategy for treating fractures in the elderly. However, the targeting proteins by UPS system in MPCs, which may contribute the decreased number of MPC in aged mice are still unknown. In present study, we used Ub-proteomics^14^ to unbiased screen Ub-proteins in MPCs and differentiated osteoblasts. We focused on pro-MPC proteins highly ubiquitinated in OB or anti-MPC proteins highly ubiquitinated in MPCs, which may be potential targets in Btz enhanced fracture repair in aged mice.

## Results

### Impaired proliferation of MPCs in aged mice

Impaired osteogenesis in aged mice can be caused either by reduced MPC proliferation, or by decreased MPC differentiation ability to osteoblasts (OBs). It is well known that aged subjects have reduced MPC and OB number in bone of aged mice^5^. However, it is not known the reduced OB number is caused by reduced MPCs number (MPC proliferation) or OB differentiation, or both. It was reported that aged people have comparable ALP level in serum^15^. To investigate whether aged mice have impaired MPC proliferation or OB differentiation, we cultured CFU-f and CFU-ALP separately in vitro using bone marrow stromal cells. Numbers of CFU-f and CFU-ALP were counted. The CFU-ALP were counter-stained the CFU-f after ALP staining. Then the ratio of CFU-ALP/CFU-f were calculated. Data showed that aged mice have reduced CFU-f and CFU-ALP, which means aged mice have reduced MPC and OB number, and reduced MPC proliferation. But the ratio of CFU-ALP/CFU-f was not changed between young and aged mice, which means MPCs in aged mice have the similar capacity to differentiate into OBs, despite their proliferation is impaired (Fig. 1A-C). To confirm the decreased proliferation of MPCs in aged mice, we immune-stained CFU-f cells with anti-BrdU and PDGFRβ antibody 4 hours after BrdU incorporation. PDGFRβ signaling plays important role in MPC proliferation (ref), which will be described below. Data showed that BrdU+ and PDGFRβ+ cells are significantly reduced in CFU-f cell of age mice (Fig. 1D). We next examined the cell proliferation and OB differentiation gene expressions in CFU-f or CFU-ALP cells, respectively. We found that all the proliferation genes, including *Ki67, CyclinD1, Cdk4* and *Pdgfrβ* are decreased in cells from aged mice, while OB differentiation genes, including *Runx2, Col-1* and *alp* are comparable in cells from aged mice. Only the late OB differentiation marker, *Ocn* was mildly increased in cells from aged mice. Data strongly supported that MPCs from aged mice have impaired proliferation in aged mice, which contribute to the decreased number of MPCs and OBs in bone of aged mice.

**Figure 1.**
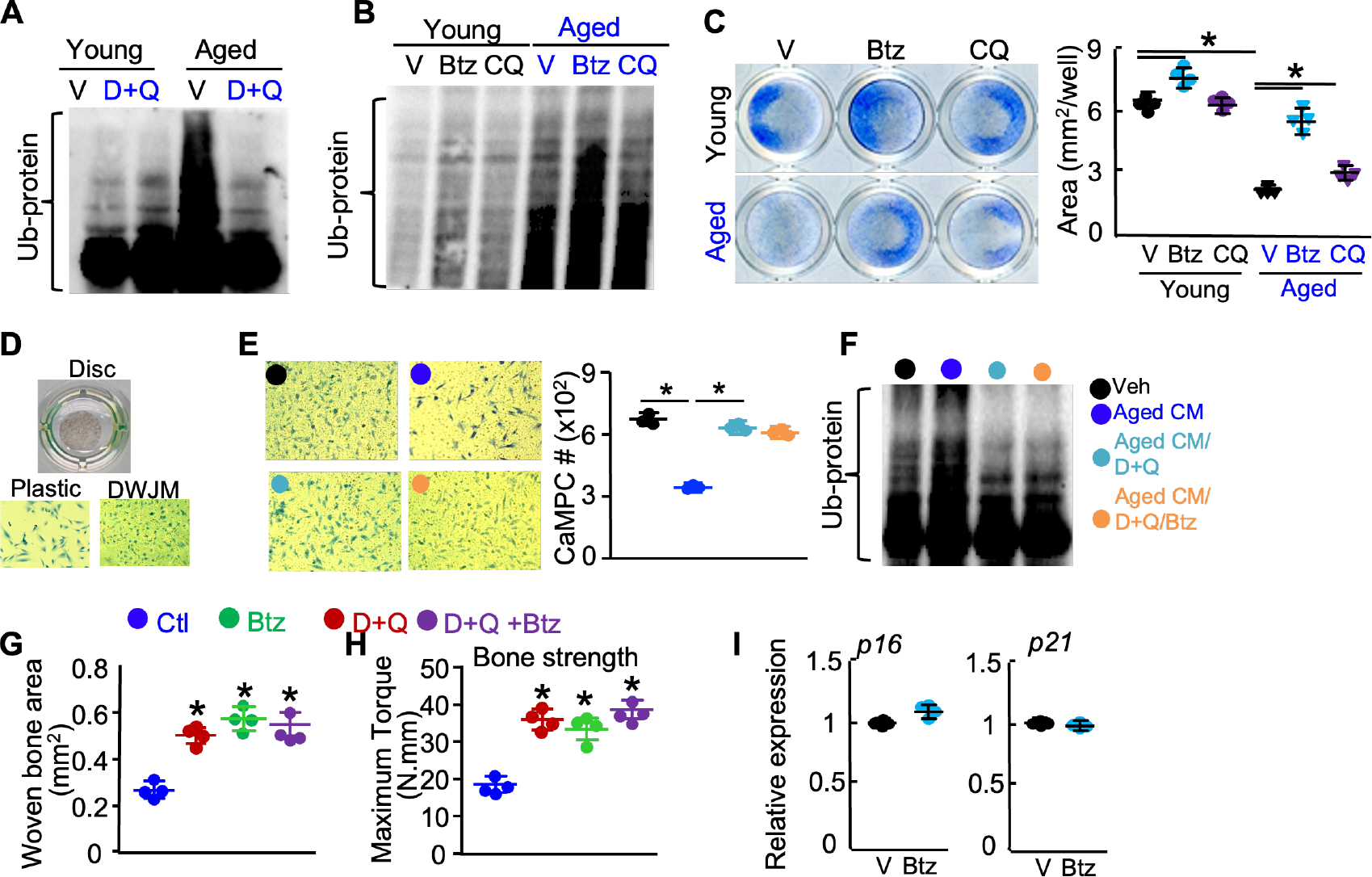
Delayed fracture healing in aged mice with decreased MPC number and angiogenesis. WT/Nestin-GFP Tg young (3m) and aged (20m) mice received an open tibia fracture and sacrificed at day 10, 14, 21 or 28. (A) Histomorphometric measurement of woven bone area in callus were performed with the Visiopharm system. N=7 mice/group. (B) microCT of bone volume in callus were performed. N=7 mice/group. (C) Biomechanical testing of tibiae at day 28. N=7 mice/group. (D) The percentage of CD45-Sca1+CD105+ MPC cells in callus at day 10 were measured by flow cytometry. (E) The Nestin+ MPC and Ub+ cells in callus at day10 were observed by fluorescent microscopy or examined by IF staining with anti-Ub antibody. Vessels in callus at d10 were examined by IF staining with anti-endomucin antibody. Values are mean ±SD. *p<0.05 vs vehicle or young group.

### Delayed fracture healing in aged mice with decreased MPC number and angiogenesis

Fracture repair was broadly reported to be delayed in aged mice^16^, we systemically examined the fracture repair in young and aged mice longitudinally. Data showed that young mice have the peak of new bone area and volume at 10 day-post-fracture (dpf), while aged mice have its peak at 14 dpf and the bone area and volume at peak is much lower than young mice (Fig. 2A%B), which infers the defect of callus formation in aged mice. Biomechanical testing at 28 dpf, the gold standard for fracture repair^17^ showed that the fractured bone in young mice recovered more than 95% of bone strength of non-fractured tibiae, while aged mice only have about 65% recovery (Fig. 2C). Bone fracture requires proper osteogenesis and angiogenesis^6^. To investigate the underlying cellular mechanism of delayed fracture repair in aged mice, we examined the number of MPC and blood vessel in callus by flow cytometry and immunostaining. Data showed that CD45-Sca1+CD105+ MPCs number examined by flow cytometry in callus at 10 dpf was significantly reduced 60% in aged mice (Fig. 2D). Nestin+ MPCs and Endomucin+ blood vessel area were reduced significantly, but the total-Ub proteins were increased in callus of aged mice (Fig. 2E).

**Fig. 2.**
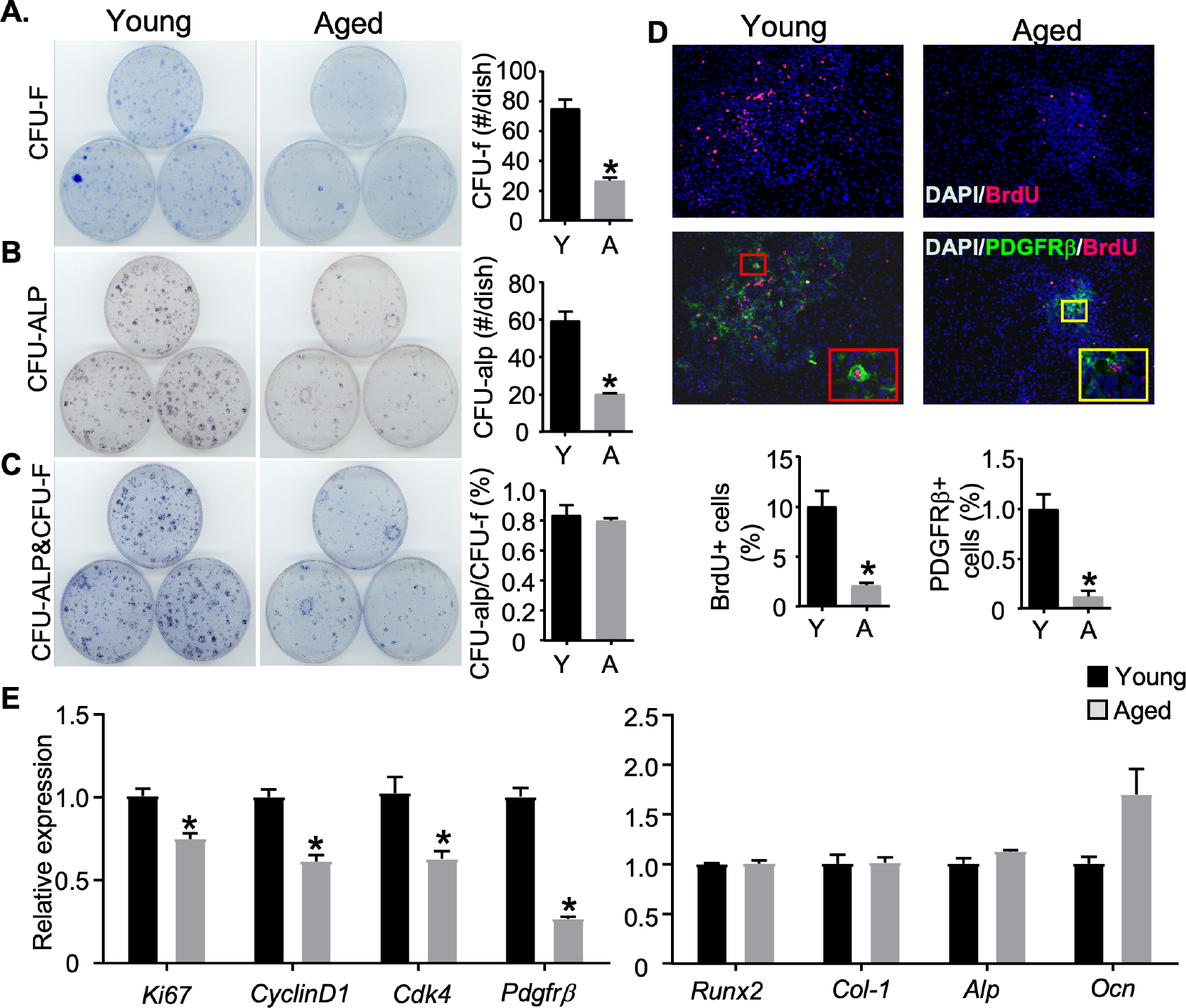
Impaired proliferation of aged BMSCs. BMSCs from Young(2m) and aged(20m) mice were cultured for 2 weeks in 15% FBS medium (A) stained with Methylene Blue for CFU-f. (B) Cells were cultured for 1 weeks in osteoblast-inducing medium and stained with 1-step NBT/BCIP reagent for CFU-alp. (C) Bone marrow stromal cells stained for CFU-ALP were stained with Methylene Blue for CFU-F. The ratio of number of CFU-alp/CFU-f. Values are mean±SEM of three dishes. (D) BMSCs from young and aged mice were examined for PDGFRβ and BrdU by IF staining. The percentage of BrdU or PDGFRβ positive cells were measured. (E) BMSCs from Young and Aged cultured w/o OB induced medium for 3d were examined for cell proliferation or OB related gene. Values are mean ± SEM of 3 mice. *p<0.05 vs young mice.

### Ub-proteomics identified PDGFRβ as the target of UPS in MPCs

We reported previously that aged mice have increased total Ub-proteins in bone and callus of aged mice^5,13^. Treatment of aged mice with proteasome inhibitor, Bortezomib (Btz) enhanced fracture repair by increasing MPC and blood vessel number in aged mice^13^. However, the targeting proteins of UPS in MPCs are still not well known, although we examined the increased expression of Runx2 and JunB in callus after Btz treatment^12^. Here, we used Ub-proteomics technology to unbiased screen the Ub-proteins in C3H10T1/2 cells and BMP2 induced OBs treated with Btz before harvesting for Ub-proteomics analysis (Fig. 3A). Because we found that MPCs have defects on proliferation, we will focus on ub-proteins that promoting cell proliferation when MPCs have lower proliferation ability (OBs), or ub-proteins that inhibiting cell proliferation when MPCs are proliferating (MPCs). Table 1 shows the top ub-proteins in both MPCs and OBs. We found that two top ub-proteins, PDGFRβ and IFITM (Fig. 3B) are associated with MPC proliferation^18,19^. We further confirmed the ubiquitination status of PDGFRβ and IFITM in MPCs and OBs by Ub assay (Fig. 3C). PDGFRβ signaling was reported to promote MPC proliferation through activating PI3K pathway and contribute to the osteogenesis and angiogenesis^20^. PDGFRβ expressing cells are required for fracture repair in adult mice^21^. Therefore, we will focus on PDGFRβ and investigate its role in regulating MPC proliferation and fracture repair.

**Fig. 3.**
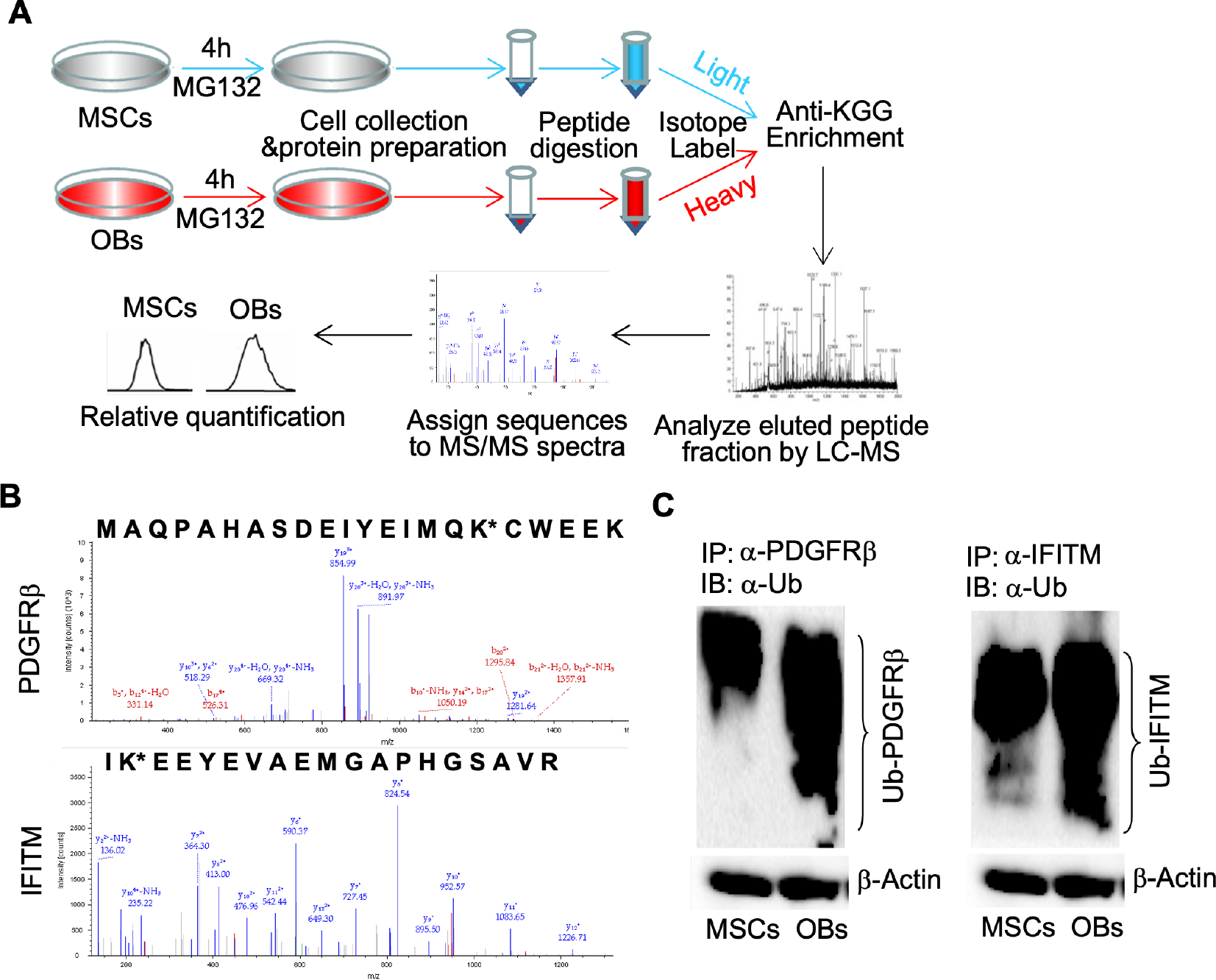
Ub proteomics identified PDGFRβ expression in aged bone callus. (A&B) C3H10T1/2 cells were treated with BMP2 (20ng/ml) for 21 days to induce OBs. MSCs and OBs cell lysates were enriched for Ub peptides followed by LC-MS/MS analysis. The flow chart of ub-proteomics (A). The representative MS/MS spectra of PDGFRβ and IFITM. (C) Ub analysis confirmed the ubiquitination in MSCs and OBs.

### Aged mice have decreased PDGFRβ in callus

To examine the role of PDGFRβ in MPCs proliferation and fracture repair in aged mice, we firstly examine the protein expression of PDGFRβ in bones. Data showed that all tissues, including bone marrow cells, cortical bone tissue and callus tissue, from aged mice have increased total ub-proteins, but decreased PDGFRβ expression (Fig. 4A). Notably, PDGFRβ protein level in callus tissue from aged mice decreased 84% compared to callus tissue from young mice, but the mRNA level of *Pdgfrβ* only decreased 50% (Fig. 4B), which means that the decreased PDGFRβ protein expression are majorly caused by the increased protein degradation. We confirmed that callus tissue from aged mice have much higher of Ub-PDGFRβ by Ub assay (Fig. 4C).

**Fig. 4.**
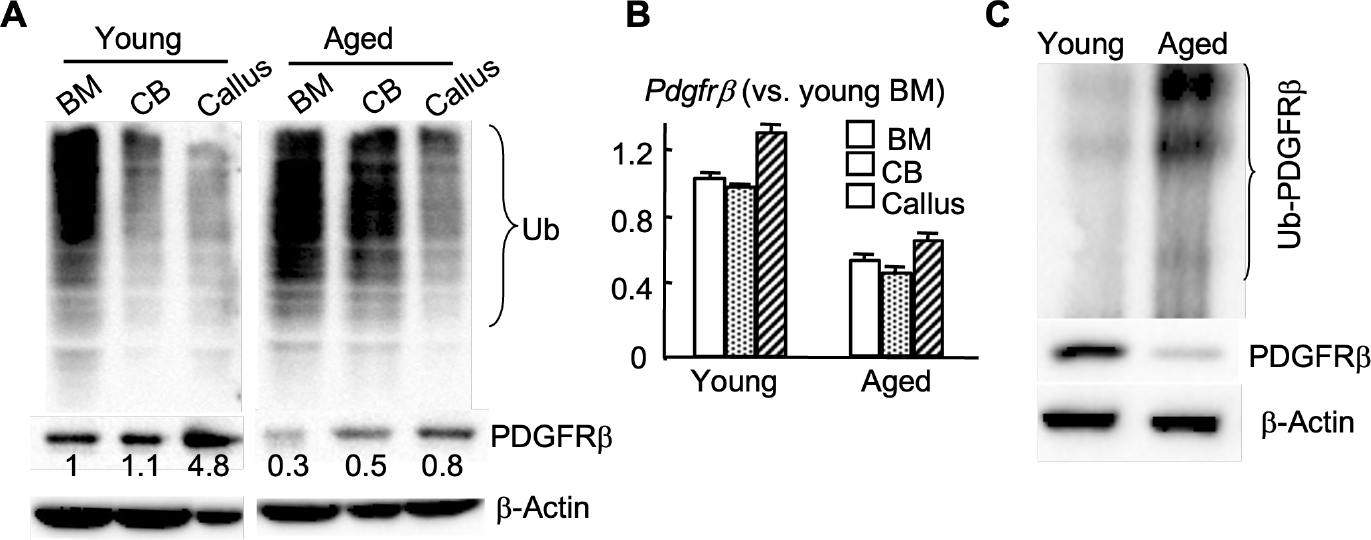
Aged mice have decreased PDGFRβ in callus. (A) Total Ub-protein and PDGFRb expressions in bone marrow (BM), cortical bone (CB) and callus tissue (d10) from young and aged mice. (B) Pdgfrβ mRNA expression. (C) Ub-PDGFRβ expression in callus tissue from young and aged mice.

### Impaired PDGF-mediated cell growth in aged mice

The ligand of PDGFRβ, PDGF was reported to strongly promote MPC growth^20^. We next examined the effect of PDGF on the growth of callus derived MPCs (CaMPCs). The CaMPCs culture method was developed recently in our lab^17^. Data showed that both PDGF and Btz alone efficiently promote the growth of CaMPCs, and they also synergistically promote even more growth of CaMPCs from young mice, which could be inhibited by PDGFRβ inhibitor. Interestingly, PDGF lost the ability to induce the growth of CaMPCs from aged mice, which was rescued by co-treatment with Btz, because Btz dramatically increased PDGFRβ expression in CaMPCs from aged mice to respond the PDGF treatment. CaMPCs with different treatment indicated were further induced into OBs and the number of ALP+ cells have the similar trend with total cells. The expression of proliferation marker, CyclinD was consistent with the cell growth results with different treatment (Fig. 5).

**Fig. 5.**
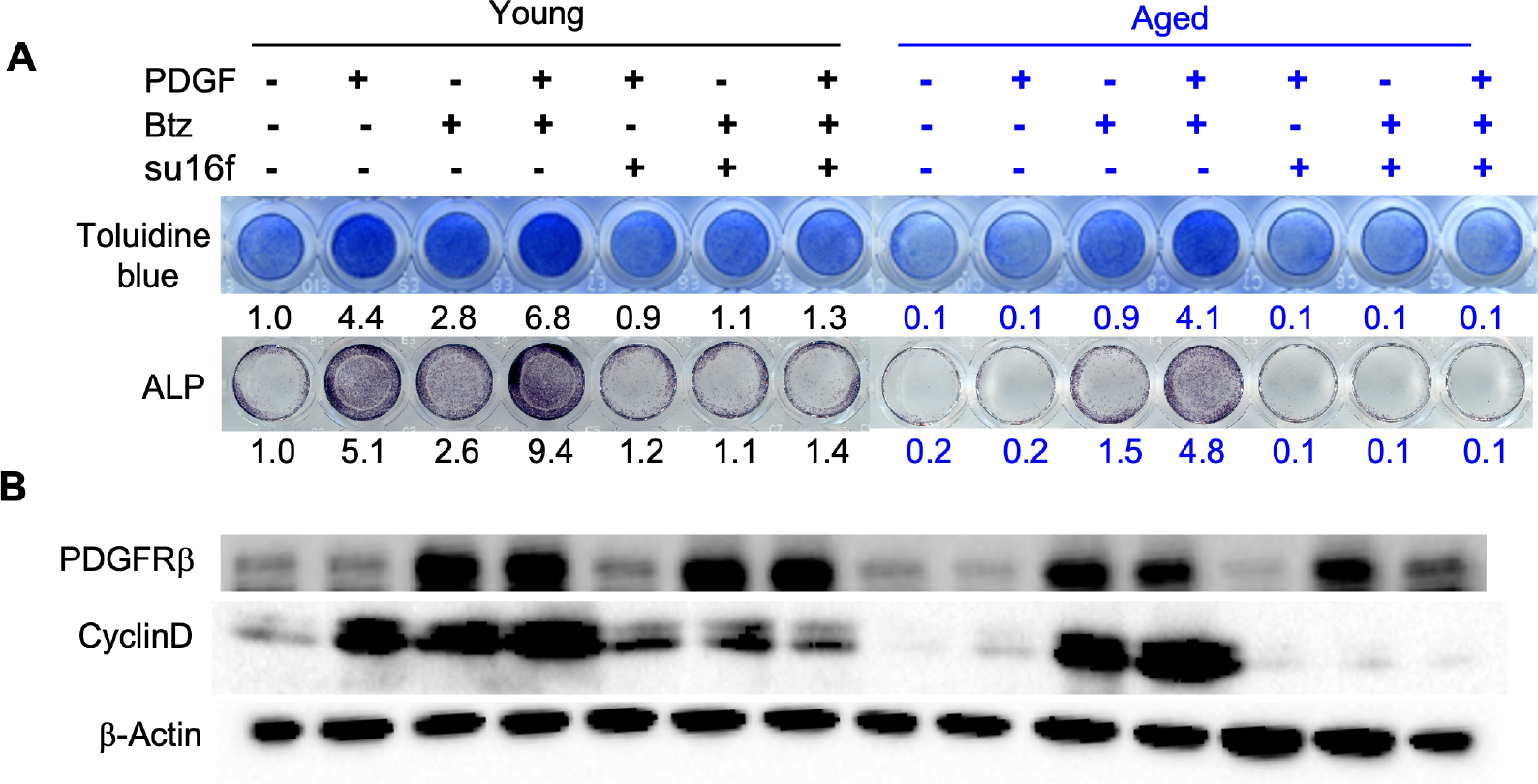
Impaired PDGF-mediated cell growth in aged mice. Callus cells isolated from young and aged WT mice at day10 post-fracture. 3^rd^ passage of cells were used. (A) callus cells were treated with 10ng/ml PDGF-BB ± 3nM Btz ±5uM su16f for 24 hours. Cells were stained with toluidine blue for cell growth. For ALP, cells were induced with OB induction medium for another 48 hours after retrieving all the treatment. Cells were stained for ALP. (B) Cells treated with 10ng/ml PDGF-BB ± 3nM Btz ±5uM su16f for 24 hours and subjected to WB for PDGFRβ and cyclinD level.

### Btz rescued the effect of PDGF on fracture repair in aged mice

We next examined the effect of PDGF on fracture repair in aged mice. As reported^12,22^, PDGF and Btz alone enhanced fracture repair in young mice evidenced by histology, microCT and biomechanical testing. Co-treatment of PDGF and Btz synergistically enhanced fracture repair. However, However, the effect of PDGF on fracture repair in aged mice is comprising, which is rescued by co-treatment with Btz, but inhibited by PDGFRβ inhibitor (Fig. 6). Our data infer that the effect of PDGF on fracture repair is depending on the expression of PDGFRβ in callus.

**Fig. 6.**
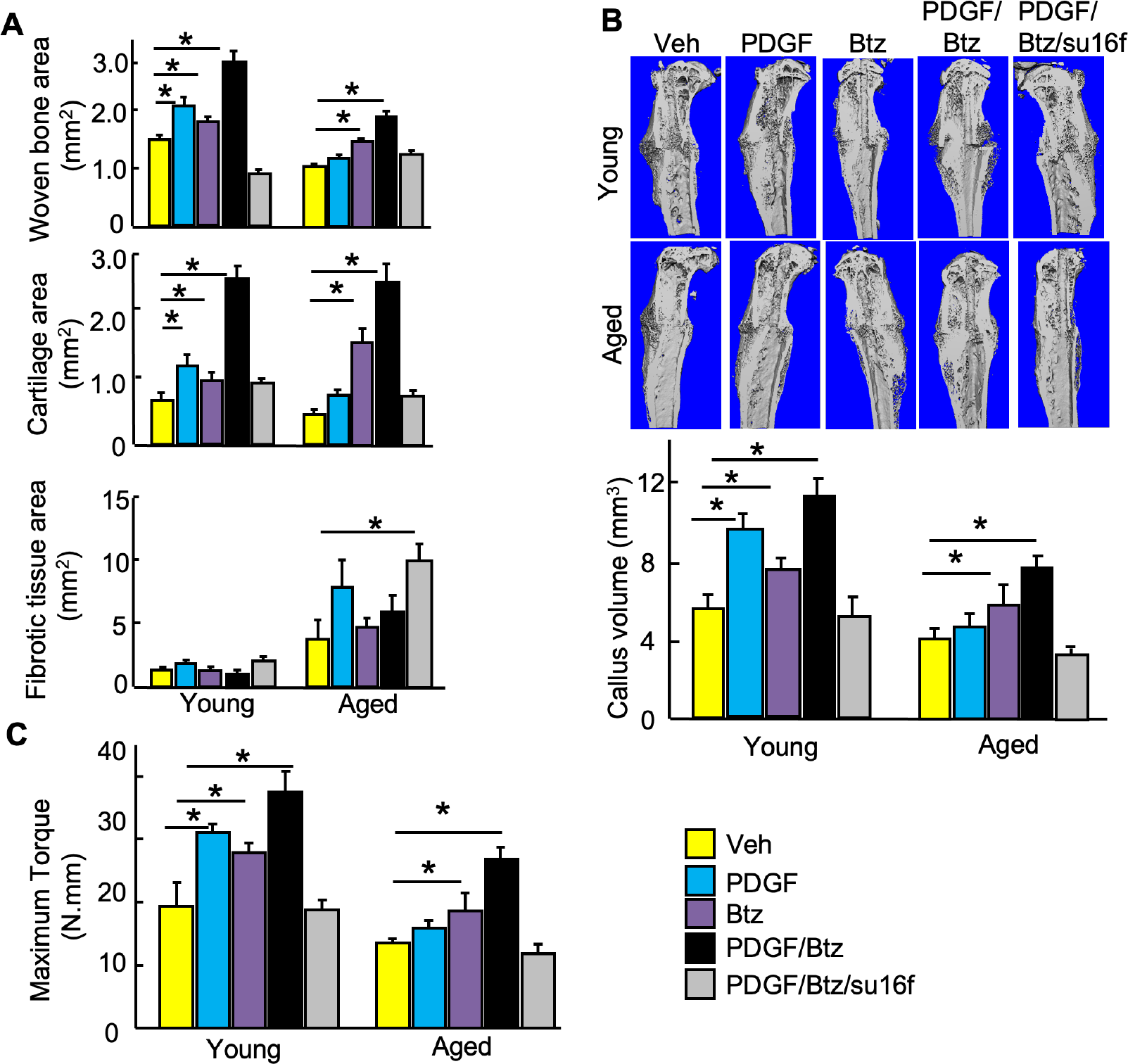
Btz/PDGF-BB synergistically enhanced bone fracture in aged mice. (A) Mice were treated with Btz as in Fig.2 Frozen sections of tibiae on d10 were stained with anti-PDGFRβ and Ub Abs. The % of PDGFRβ+ cells in callus were measured. Values are mean±SD of 3 mice. Mice were treated with PDGF-BB (1.5ug/local injection) ± Btz (0.6mg/kg by i.p.) ± su16f (10mg/kg by gavage) as Fig2. (B) Histomorphometric measurement of woven bone area, cartilage area and fibrotic tissue area in callus at day 14 after fracture were performed with the Visiopharm system. (C) microCT was performed to measure the callus volume. (D) Biomechanical testing was performed on tibiae at day 28. Values are mean±SD of 7 mice.

### Temporal depletion of PDGFRβ+ cells impaired fracture repair in young mice

To examine the effect of decreased PDGFRβ+ cell number on fracture repair in aged mice, we used Pdgfrb^rtTA^;tetO-DTA mice to temporal depletion of PDGFRβ+ cells at MPC proliferating stage and mimic the microenvironment in callus of aged mice (Fig. 7A). Data showed that single dose of Dox treatment to activate the rtTA-tetO system significantly reduced PDGFRβ+ MPCs and endomucin+ blood vessels in callus at 10 dpf and start to recover at 14 dpf (Fig. 7B) in young mice. PDGFRβ+ cells depletion by Dox treatment significantly impaired fracture repair in young mice evidenced by woven bone, cartilage area and biomechanical testing, but has no significant effect in aged mice (Fig. 7C&D). Our data infer that decreased number of PDGFRβ+ cells was the cause of impaired fracture repair in aged mice.

**Fig. 7.**
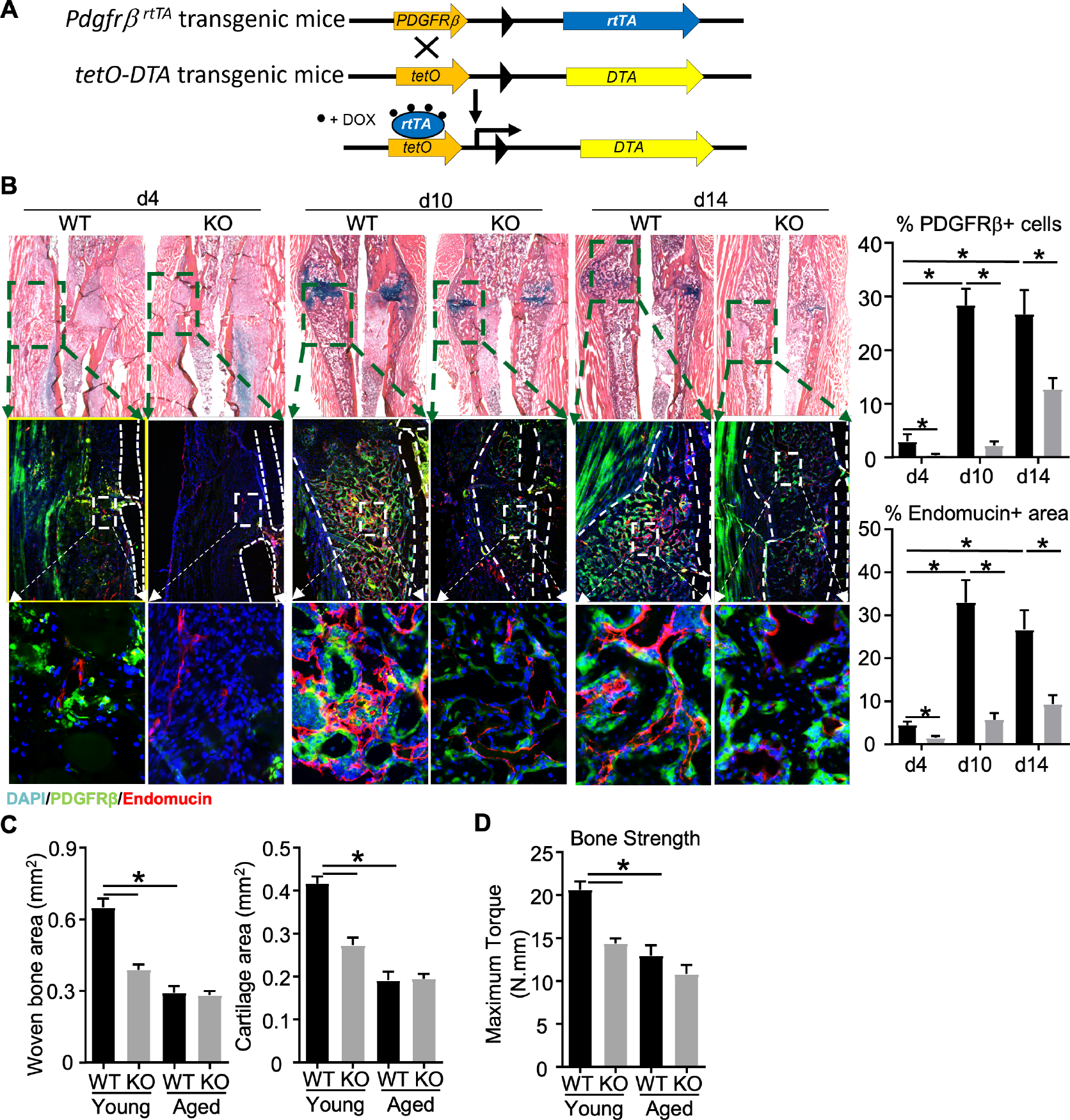
Temporal knockout of PDGFRβ+ cells impaired fracture repair in young mice. (A) Generation of *Pdgfrβ-rtTA/TetO-DTA* mice. (B) 3-month-old mice were treated with Dox 1 day before fracture surgery. Frozen sections of tibiae on d4, 10 and 14 were stained with anti-PDGFRβ and Endomucin Abs. The % of PDGFRβ+ cells and Endomucin+ area in callus were measured. Values are mean±SD of 3 mice. (C) Histomorphometric measurement of woven bone area and cartilage area in callus at day 10 after fracture were performed with the Visiopharm system. (D) Biomechanical testing was performed on tibiae at day 28. Values are mean±SD of 7 mice.

## Discussion

Proteasomal protein degradation is one of major protein degradation pathways and is mediated by the UPS^23^. Proteasomal degradation of intracellular proteins, including transcription factor or abnormal proteins trafficking from the cytosol nuclei, plays an important role in bone cell regulation under normal and pathological conditions. A number of positive regulators OB differentiation, including the downstream of BMP or TGFβ signal, Smads, Runx, JunB, for were reported to tightly regulated by UPS^2-5^. Inhibition of UPS system by proteasome inhibitors prevents ovariectomy-induced osteoporosis and promote in vivo bone regeneration by stabilizing Runx2 in MPCs^10^. Local delivery of proteasome inhibitor enhanced fracture in rats^24^. Our lab discovered the temporal increase of total ub-proteins and multiple E3 ligases, including Wwp1, Itch and Smurf2 at early stage of fracture repair^12^, which provides the rationale of treating fracture with proteasome inhibitor only at early stage, because long time treatment of proteasome inhibitor influences osteoclast formation at remodeling stage of fractur repair. Long-term use of proteasome inhibitors inhibits osteoclast differentiation by inhibiting NF-κB signaling^25^. Using Nestin-GFP MPC reporter mice, we found proteasomal inhibition not only promoted OB differentiation^10^(18219387), but also increased number of MPC in fracture repair^12^.

Fracture repair in adult young mice can be perfectly healed by themselves, which reduced the significance of the findings of proteasome inhibitor on fracture healing. Aged mice have been reported to have delayed fracture repair^16^. In clinic, elderly people with fracture have a high incidence of disability and mortality. However, there is still no FDA approved drug for treating fracture for elderly people. Thus, there is a unmet need to develop new therapy for fracture repair in elderly people. We reported previously that MPCs in MPCs from aged mice have increased ubiquitinated JunB and Runx2^4,5^ and dramatic increase of total Ub-protein at early stage in callus of aged mice^13^. Present study also found that the major defect of MPCs is decreased MPC proliferation resulting in decreased number of MPCs in bone marrow, but surprisingly MPCs from aged mice have comparable OB differentiation ability. These findings provide the basis of treating aged mice with proteasome inhibitor. Eventually, we reported that proteasome inhibitor also enhanced fracture repair in aged mice with increased MPCs number after treatment^13^.

Although fantastic results of proteasome inhibition were achieved in treating fracture repair both in young and aged mice, the mechanism of increased MPC number by proteasome inhibitors is unknown. Ub-proteomics technology^14^ provide the chance to investigate the targets of proteasome inhibitors in MPC potentially regulating MPC proliferation. Our results showed the top ubiquitinated proteins both in MPCs and OBs. We focused on those Ub-proteins with anti-MPC in MPCs and pro-MPC in OBs, when MPCs lost the proliferation ability by degrading pro-MPC proteins. Among the top Ub-proteins, PDGFRβ is eye-catching, because PDGFRβ plays important role in regulating both osteogenesis and angiogenesis, which were observed decreased in callus of aged mice, although another target, IFITM is also associated with MPC proliferation^18^. Our data showed a significant decrease of PDGFRβ at protein level and increase of Ub-PDGFRβ, but mild decrease at mRNA level, suggesting the dys-regulated protein post-translational modification majorly contributes to the decreased expression of PDGFRβ. PDGFRβ+ cells are reported to be required for fracture healing in adult mice^21^ and PDGFRβ signaling is critical for MPC proliferation and fracture repair in adult mice^12^. Notably, MPCs derived from callus of aged mice and aged mice with fracture did not respond to PDGF-BB treatment anymore. These findings are consistent with previous reports by others. PDGF containing platelet-rich plasma failed to heal the bone defect in aged sheep^26^. Platelet-derived wound healing factors failed to healing skin wound in patients with the age between 57-75^27^ compared to a controversial study with excellent therapy effect on young patients with the age under 57 (50% < 45 years)^28-30^. There is a difference on PDGF-BB induced cell migration between young and aged endothelial cells^31^. Our data explained the phenomena discovered by our colleagues and provide a solution. Proteasome inhibitor, Btz restored the ability of PDGF-BB on cell proliferation and fracture healing caused by decreased PDGFRβ expression.

Collectively, the present study not only identified the protein target of proteasome inhibitor in MPCs, but also connected the compromised effect of PDGF treatment on diseases with PDGFRβ proteasomal degradation. We provide an innovative therapy strategy with the combination of PDGF-BB or its product with proteasome inhibitor in clinic.

## Materials and methods

### Animals

Young (3-month-old, 26-year-old in human) and aged (20-month-old, 62-year-old in human) C57BL/6J (WT) mice from National Institution of Aging were used. Mice were housed in micro-isolator technique rodent rooms. *Nestin-GFP* reporter mice that were originally generated in Dr. Tatyana V Michurina (Cold Spring Harbor Laboratory)^32^ and breeded on a C57BL/6J background, in which the 5.8-kb fragment of the promoter region was inserted upstream of sequences encoding destabilized eGFP as we described previously^12^. Pdgfrβ^rtTA^/TetO-DTA mice were crossed by Pdgfrβ^rtTA^ (Strain #:028570) and TetO-DTA (Strain #:008168) mice, which are purchased from Jax. Mice were given Doxycyclin (Dox) once right before fracture surgery by i.p. (5mg/kg). All animal procedures were approved by the University Committee on Animal Research at the University of Rochester.

### Tibial fracture procedure and animal treatments

Open tibial fracture procedures were performed according to standard operation procedure established in Center for Musculoskeletal Research6. In brief, an incision of 5 mm in length was made in the skin over the anterior side of the tibia after anesthesia. A sterile 27 G x 1.25-inch needle was inserted into the bone marrow cavity of the tibia from the proximal end, temporarily withdrawn to facilitate transection of the tibia using a scalpel at midshaft, and then reinserted to stabilize the fracture. The incision was closed with 5-0 nylon sutures. Fractures were confirmed by radiograph. Callus tissues were harvested on day 10, the time when soft callus is formed, following fracture procedure for cell preparation. Mice received slow-released extended release (XR) buprenorphine, 0.5 mg/kg, to control pain. Fractured mice were given Btz (Bortezomib, 0.6mg/kg by i.p.), su16f (PDGFRβ inhibitor 10mg/kg by gavage), PDGF-BB(1 μg/10 μL, R&D, by local injection) or vehicle at 1, 3, 5, and 7 dpf.

### Preparation of CaMPCs

Callus derived mesenchymal progenitor cells (CaMPCs) are obtained as described previously^33^. Briefly, callus tissues were dissected from the tibia, cut into small pieces (<1 mm3), and digested in 1 ml of Accumax solution (STEMCELL, 1 hour, room temperature). Cells were passed through a 35 um-filter and red blood cells were lysed with ammonium chloride (5 minutes, room temperature). Cells were cultured in basal medium (alpha-MEM medium containing 15% FBS). Cells that migrated from callus pieces were cultured in the basal medium to confluence, and CaMPCs in the 3^rd^-5^th^ passage were used for experiments.

### Flow cytometry

APC-anti-CD45, PE-anti-CD105 and PE-cy7-anti-Sca1 antibodies (Abs) are purchased from eBioscience. Callus and BM cells are stained with various fluorescein-labeled Abs and subjected to flow cytometric analysis using a Becton-Dickinson FACSCanto II Cytometer, according to the manufacturer’s instructions. Results are analyzed by Flowjo7 data analysis software (FLOWJO, LLC Ashland, OR).

### Histology and histomorphometric analysis

Tibiae were fixed in 10% formalin and decalcified in 10% EDTA. Paraffin sections were prepared for histology, and 4 μm thick sections were cut at 3 levels (each level was cut 50 μm apart). Sections were stained with Alcian blue/hematoxylin (ABH) and counterstained with eosin/Orange G, following by scanning with an Olympus VS-120 whole-slide imaging system. Scanned images were labeled by numbers, and histomorphometric measurement of cartilage and woven bone area was performed using Visiopharm software (version 2018.4) following CMSR SOPs in a blinded manner.

### Immunostaining and data analysis

Tibiae were fixed in 10% formalin for 48 hours at 4°C, decalcified in 14% EDTA for two weeks, processed, and embedded in optimal cutting temperature compound (Tissue-Tek). Seven-micron sagittal sections were cut and probed with antibodies for Ub (Santa Cruz, cat#: sc-8017), Endomucin (Santa Cruz, cat#: sc-65495) or PDGFRβ (Santa Cruz, cat#: sc-80991), followed by secondary antibodies and counterstained with Dapi. The Sections were imaged with a Zeiss AxioImager motorized fluorescent microscope system with AxioCam camera. The percentage of PDGFRβ^+^, Endomucin^+^, and Ub^+^ area within callus was quantified with ImageProPlus software.

### Micro-CT and Bio-mechanical testing

For microCT, fractured tibiae at 10 dpf were dissected free of soft tissue, fixed overnight in 70% ethanol and removed the stabilizing needle before scanning at high resolution (10.5 μm) on a VivaCT40 micro-CT scanner (Scanco Medical, Basserdorf, Switzerland) using 300 ms integration time, 55kVp energy, and 145 uA intensity. 3-D images were generated using a constant threshold of 275 for all samples. For Bio-mechanical testing, fractured tibiae at 28 dpf are stored at −80°C after removing the stabilizing needle carefully to avoid any damage to the architecture of the callus. The tibial ends are embedded in polymethylmethacrylate and placed on an EnduraTec system (Bose Corporation). A rotation rate of 1°/s is used to twist the samples to failure or up to 80°. Maximum torque, maximum rotation, and torsion rigidity are analyzed using a CMSR SOP^12,13,17^.

### Quantitative real time qPCR

Callus tissues or cultured MPCs were subjected to RNA extraction with TRIzol, and cDNA was synthesized using the iSCRIPT cDNA Synthesis kit (BioRad). qPCR was performed with iQ SYBR Green Supermix using an iCycler PCR machine (BioRad). The fold change of gene expression was first normalized to actin and then normalized to the values in control group.

### Ub-proteomics assay

C3H10T1/2 cells were cultured and labeled with ^13^C_6_^15^ N4-arginine and ^13^C6-lysine for five passages. For OB differentiation, C3H10T1/2 cells were induced by BMP2 (20ng/ml) for 21 days. Differentiated cells are not labeled. Both cells are treated with MG132 for 4 hours before harvest. Proteins are qualified after extraction. Equal amount of proteins from undifferentiated and differentiated cells were mixed and send to our Proteomics Core in University of Rochester. The following proteomics was completed in the proteomics core as described in the publication^14^. Briefly, proteins are carboxamidomethylated and digested with trypsin. After digestion, peptides were purified for the K-GG modification using the K-GG motif antibody kit (Cell Signaling). Eluted peptides were desalted and concentrated with PerfectPure C18 tips following with LC-MS/MS analysis. Tandem mass spectra were collected with an LTQ Orbitrap hybrid mass spectrometer.

### Ubiquitination assay and western blot analysis

For ubiquitination assay^34^, MPCs were treated with the proteasome inhibitor MG 132 (10 μM) for 4 hours. Whole-cell lysates (200 μg protein/sample) were incubated with UbiQapture-Q Matrix (Biomol) by gentle agitation at 4°C overnight to pull down all ubiquitinated proteins according to the manufacturer’s instructions. After washing three times, captured proteins were eluted with 2× SDS-PAGE loading buffer and analyzed by Western blotting using anti-PDGFRβ antibody, as described previously (ref). For western blot analysis, callus tissues homogenized under liquid nitrogen or CaMPCs were extracted proteins with RIPA lysis buffer. Proteins were quantitated using a kit from Bio-Rad and loaded onto 10% SDS-PAGE gels and blotted with anti–Ub, PDGFRβ, IL-1, CyclinD1 or actin antibodies. Bands were visualized using ECL chemiluminescence (Bio-Rad, catalog 1705061).

### Statistical analysis

Statistical analysis was performed using GraphPad Prism 5 software (GraphPad Software Inc., San Diego, CA, USA). Comparisons between two groups were analyzed using a 2-tailed unpaired Student’s t-test. One-way ANOVA and Tukey post-hoc multiple comparisons were used for comparisons among three or more groups.

